# Age-specific survival in an English Twite population

**DOI:** 10.1101/2021.11.24.469575

**Authors:** Ismini Gkourtsouli-Antoniadou, Steven R. Ewing, George Hudson, Michael A. Pearson, Julia Schroeder, Peter E. Welch, Nicholas Wilkinson, Jamie Dunning

## Abstract

Like many bird species associated with agricultural habitats in the UK, the Twite *Linaria flavirostris* has undergone severe declines over recent decades due to habitat degradation, with populations in England, Wales and Ireland now restricted to a few small pockets. However, the demographic drivers of these declines are still largely unresolved. We estimated the survival of Twite from a small population at the southernmost edge of the English range in Derbyshire using capture-mark-recapture data from 2016–2019. Annual apparent survival for juveniles (0.14-0.34) was lower than for adults (0.29-0.56) and less than that of other Cardueline finches. Our results suggest that low juvenile survival may be one demographic driver underpinning the recent decline of the Derbyshire Twite population, although we also cannot rule out the possibility that differences in emigration of juveniles and adults from the population also contribute to the observed age-specific apparent survival rates.

## Introduction

Twite *Linaria flavirostris* are a cardueline finch, with a disjunct population, split between northwest Europe and central Asia (Perrins 1987). In the UK, Twite breed in areas of low-intensity upland agriculture, managed moorland edge and rocky coasts, and winter in seed-rich farmland and salt marsh (Raine 2006). Twite are a species of conservation concern in the UK (Eaton *et al.* 2015). Periodic national surveys show that the population declined by 21% between 1999 and 2013, from an estimated population size of 9948 (95% CI 6293-14734) to 7,831 pairs (95% CIs: 5829-10137; Langston *et al.* 2006, Wilkinson *et al.* 2018). These declines are likely underpinned by the intensification of farming practices in both the breeding and wintering areas (Wilkinson & Wilson 2010), but factors driving the decline are still unresolved.

The remnant breeding distribution of Twite in England is heavily fragmented (Dunning *et al.* 2020), occurring in isolated sub-populations (hereafter, populations) in the South Pennines and North-West Derbyshire (Dunning *et al.* 2016), which together represent the core English breeding range. The South Pennine population declined by 72% between 1999-2013 (Wilkinson et al. 2018), and may be as few as c. 30 breeding pairs in 2021, despite recent conservation interventions to restore upland meadow pasture (Wilkinson *et al.* 2018; Wilkinson *et al.* in prep.). The Derbyshire population, which was once the southernmost extent of the Pennine range, has not been monitored as comprehensively, but local records indicate a parallel decline in this population since the 1970s (Orford 1973, DOS 2013).

Demographic drivers of the Twite’s population decline in England are unclear. Comparisons of Twite demographic rates demonstrated that both whole-season breeding productivity and survival were lower in the small South Pennine populations than larger populations on the Western Isles of Scotland (Wilkinson *et al.* in prep.), providing some insight into potential drivers of population change in England. However, where survival estimates have been derived for Twite in the past, ringing data were limited and recapture rates low (Raine *et al.* 2006; Wilkinson *et al.* in prep), compromising the accuracy of survival estimates.

Here, we add to the current understanding of the demographic parameters potentially underpinning the species’ population decline by estimating survival for Twite breeding in North-West Derbyshire between 2016-2019, where they have been the subject of an intensive capture-mark-recapture study. We investigated whether there was evidence for time-dependent survival and differences between age groups in this isolated population. We then compared our findings with other published and unpublished estimates of survival as a basis for examining plausible demographic drivers of population declines in Twite. Our results offer useful insights into the survival of Twite breeding in England and provide evidence for targeted conservation interventions for this species.

## Methods

We monitored a Twite population near the village of Dove holes in North-West Derbyshire (SK0878), close to the Peak District National Park, between 2016 - 2019. Land use in this area consists primarily of sheep-grazed pastures and limestone quarrying. Twite mainly nested in cavities on limestone cliff faces (Frost 2008), making access for monitoring prior to fledging impossible. Birds were instead captured at a baited feeding station (see Raine *et al.* 2009, Dunning *et al.* 2016) using whoosh nets during two phases: first during the pre-breeding period in spring (March - April), when birds returned from their winter grounds in the South-East of England; and then, in the post-breeding period in late summer and autumn (July - October).

We fitted each bird with a unique combination of a British Trust for Ornithology (BTO) metal ring and three plain colour rings. We aged birds following standardised EUring codes (Du Feu *et al.* 2020) as either juveniles or adults based on plumage and moult characteristics. However, although we were able to sex some adults on behaviour, genetics or extreme biometrics, most birds could not be sexed on plumage alone at the point of capture due to unreliable sexing criteria, particularly pink rumps indicating male birds (Svenson 1992, McLoughlin *et al.* 2012). Systematic resightings of colour-marked birds at the feeding station were made by two observers every other day from March to October, for intervals of one to three hours at a time. We also included opportunistic resightings from experienced birdwatchers, where they could be verified with either a clear photograph or through multiple, corroborated observations. This intensive resighting effort led to very high detection rates of marked birds.

### Survival analysis

We used mark-recapture modelling approaches implemented via programme MARK (White & Burnham 1999) and the RMark package (Laake 2013) in R (version 3.6.1; R Core Team 2021) to estimate annual survival probabilities for Twite in Derbyshire. Twite were marked and resighted in this study over six months, which potentially violates the instantaneous sampling assumption of mark-recapture analysis. To counter this, we only used birds captured and resighted as juveniles or adults during August, September, and October between 2016-2019. We used data from these three months as it is immediately post-fledging, and thus covers the period when all Twite have been marked in our study system. It is also prior to the migration period, thus reducing the possibility that we sampled birds breeding outside of the study population. We considered the possibility of juvenile dispersal from the study site described by Raine et al. (2006) to be low in this population due to its isolation from the remainder of the core English breeding range. For each Twite marked between August-October, we created an annual resighting history with 4 capture occasions (2016-17; 17-18; 18-19; 19-20) based on their initial capture date and subsequent resightings in the study area.

We employed a capture-mark-recapture model modelling framework to calculate apparent survival probability (phi) and resighting probability (p; Lebreton *et al.* 1992). We started by fitting simple survival models that assumed constant survival and resighting probability and then developed more elaborate parameterisations of both. We fitted different models assuming age- and time-dependence in resighting probability. Some of these models returned warning messages suggesting that not all detection probability parameters were estimable. This was simply due to high resighting effort yielding age categories and time periods with detection probabilities equal to 1, and thus we proceeded with consideration of these models . We then fitted models specifying age-dependent survival, with individuals coded either as juveniles (age code three when initially captured), or adults (classified as age codes four, five and six). Individuals were only classified as juveniles during their first year and thereafter graduated to the adult age category for subsequent years. Birds of unknown age were dropped from the analyses. We also examined whether survival of Twite varied between years.

We used Akaike’s Information Criterion corrected for small sample sizes (AICc) to assess the relative support for different competing models of survival (Burnham & Anderson 1998). The models were ranked by their AICc values and the most parsimonious model (the model with the highest explanatory power and minimum parameters) was the one with the lowest AICc (Grosbois *et al.* 2008). Furthermore, to examine the extent to which the Twite mark-resighting data were overdispersed, we used Fletcher’s ĉ as provided by the output of the global (i.e. phi(age+time) p(age+time)) model, as other tests such as RELEASE were not an option due to data sparsity. Where data are overdispersed (ĉ > 1), it is generally advised that an adjusted version of AICc, specifically QAICc, is used for model selection.

## Results

Between 2016-2019, we captured 229 individual Twite, of which 165 (72.0%) were ringed as juveniles, 52 (22.7%) as adults and 12 (5.2%) were of unknown age. Both the numbers of birds resighted, and juveniles ringed declined each year despite high capture success and resighting effort throughout the study (Table 1).

**Table 1.**
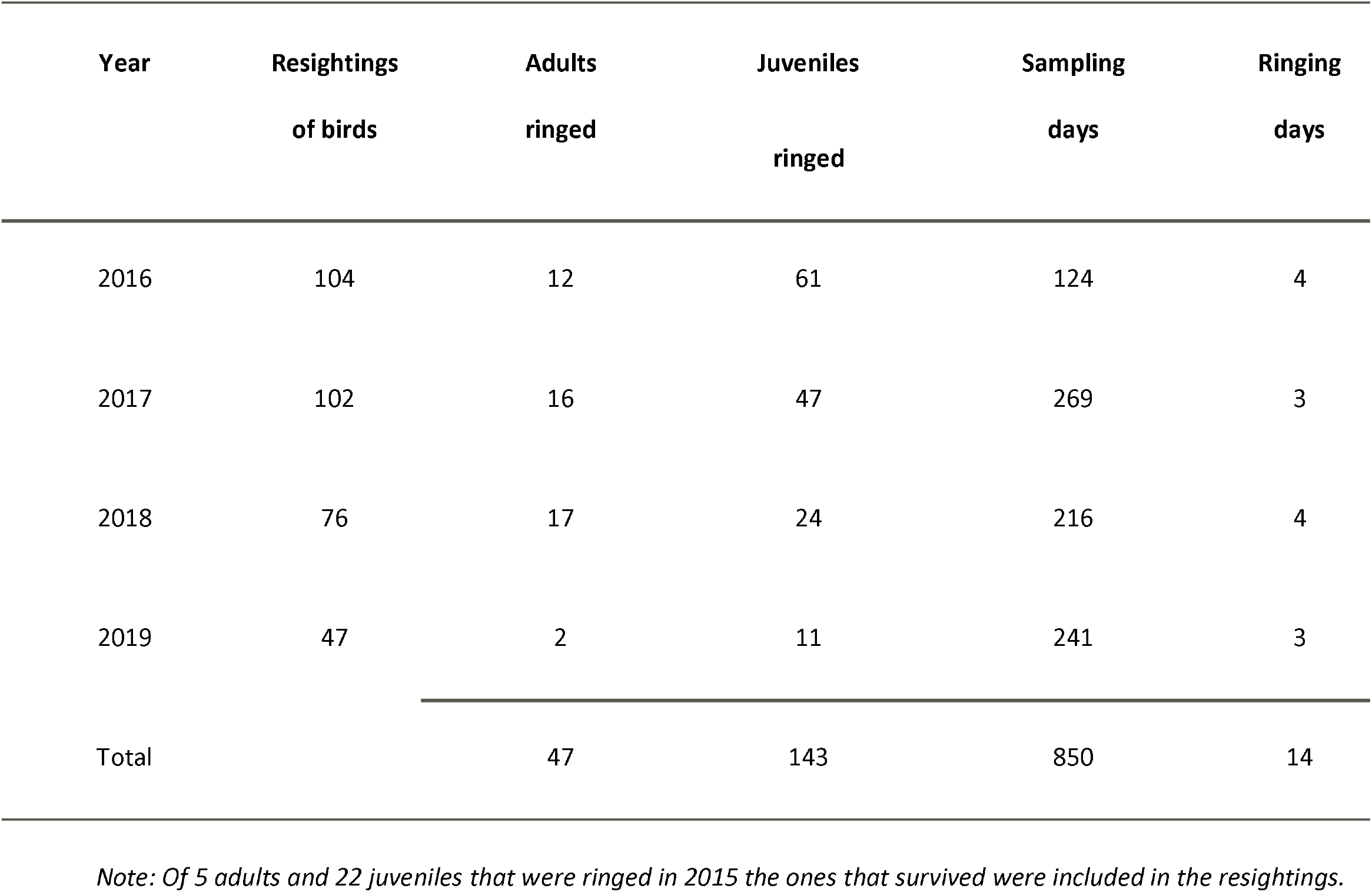
Numbers of individually marked Twite resighted, juveniles and adults ringed, and resighting and ringing effort (days) in N.W. Derbyshire each year from March to October, excluding birds of unknown age.

| **Year** | **Resightings**<br>**of birds** | **Adults**<br>**ringed** | **Juveniles**<br>**ringed** | **Sampling**<br>**days** | **Ringing**<br>**days** |
| --- | --- | --- | --- | --- | --- |
| 2016 | 104 | 12 | 61 | 124 | 4 |
| 2017 | 102 | 16 | 47 | 269 | 3 |
| 2018 | 76 | 17 | 24 | 216 | 4 |
| 2019 | 47 | 2 | 11 | 241 | 3 |
| Total |  | 47 | 143 | 850 | 14 |

*Note: Of 5 adults and 22 juveniles that were ringed in 2015 the ones that survived were included in the resightings.*

### Annual apparent survival

The estimate of overdispersion (ĉ) for the global model was 1.08 and AICc was deemed suitable for model selection. The best-supported mark-recapture model assumed that apparent annual survival varied as a function of time and an additive effect of age, with constant resighting probability (model 1, Table 2). The next best model only included age, but not time-dependence, in survival, but this had an AICc value 2.6 points greater than the top model (Table 2), indicating less support. According to the top model, the resighting probability of marked Twite was 0.86 (95% CIs: 0.58 - 0.96). This model also showed that survival was higher for adults than juveniles, and varied between years, although temporal variation was similar between the two age classes. Adult and juvenile survival ranged between 0.29-0.56 and 0.14-0.34 respectively (Fig.1, Table 3).

**Table 2.** Comparison of CJS models for estimating the survival probability of Twite. [Annual survival probability=Phi, recapture probability=p, number of estimable parameters=K, Akaike’s information criterion values=AICc, AICc differences=Δi/DeltaAICc, AICc Weights=W_i_, constant

| No. | Model parameters ($\Phi$)($p$) | K | AICc | DeltaAICc | Wi |
| --- | --- | --- | --- | --- | --- |
| 1 | $\Phi(\text{age}+\text{time})p(c)$ | 5 | 257.49 | 0 | 0.59 |
| 2 | $\Phi(\text{age})p(c)$ | 3 | 259.99 | 2.5 | 0.17 |
| 3 | $\Phi(\text{age}*\text{time})p(c)$ | 7 | 260.83 | 3.34 | 0.11 |
| 4 | $\Phi(\text{age})p(\text{age})$ | 4 | 262.07 | 4.58 | 0.06 |

estimate= $c$ ].
| 5 | phi(time)p(c) | 4 | 262.36 | 4.87 | 0.05 |
| --- | --- | --- | --- | --- | --- |
| 6 | phi(c)p(time) |  | 266.31 | 8.82 | 0.00 |
| 7 | phi(c)p(c) | 2 | 266.73 | 9.24 | 0.00 |
| 8 | phi(c)p(age) | 3 | 268.79 | 11.3 | 0.00 |
| Species | Study period | Adult annual | Juvenile annual | Reference | Population |
|  |  | survival (SE) | survival (SE) |  | trend in the |

**Fig.1.**
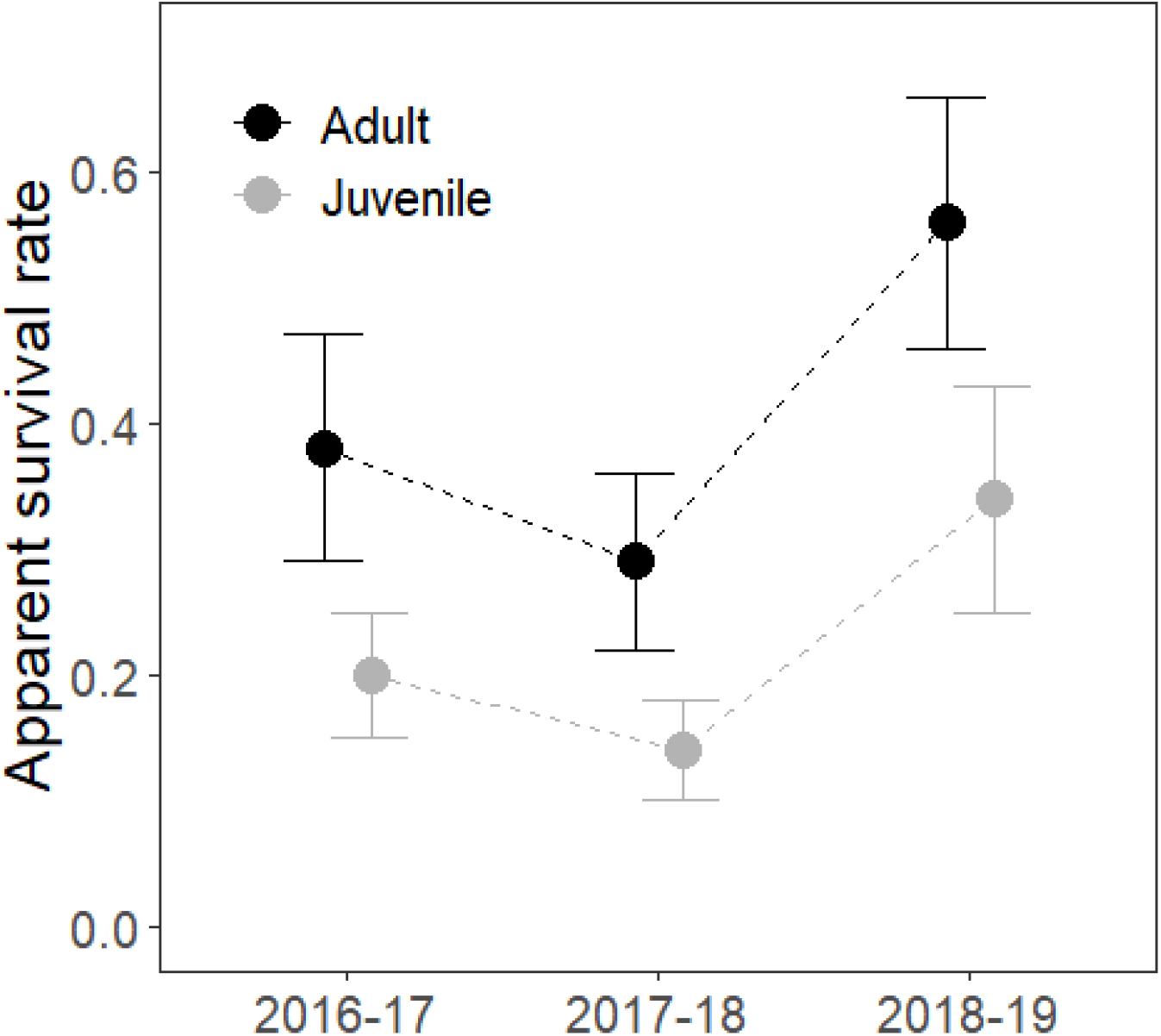
Apparent survival for the adult and juvenile Twite between each year (2016-2019).

**Table 3.** Annual adult and juvenile survival estimates including standard errors (SE) for the Twite and other Cardueline finches with similar habitat preferences in the UK.

|  |  |  |  |  |  |
| --- | --- | --- | --- | --- | --- |
| Twite in Derbyshire | 2016-17 | 0.38 (0.09) | 0.20 (0.05) | current study | Declining |
|  | 2017-18 | 0.29 (0.07) | 0.14 (0.04) |  |  |
|  | 2018-19 | 0.56 (0.10) | 0.34 (0.09) |  |  |
| Twite in S. Pennines | 2003-10 | 0.24-0.44 | 0.24-0.44 | Wilkinson *et al.* in prep. | Declining |
| Twite in N.W. Scotland | 2007-19 | 0.35 males | — | BTO | — |
|  |  | 0.33 females |  |  |  |
| Linnet | 1962-95 | 0.37 (0.02) | 0.34 (0.03) | Siriwardena *et al.* 1998 | Declining |
| Goldfinch | 1962-95 | 0.37 (0.02) | 0.34 (0.02) | Siriwardena *et al.* 1998 | Declining |

## Discussion

Apparent survival of first-year Twite in the Derbyshire study population was lower than that of adults. As these individuals are likely not mixing with other flocks during the breeding season (Dunning *et al.* 2016), exhibit high site fidelity (Raine 2006) and are isolated from the South Pennine populations, we suggest that it is more likely that the lower apparent survival rates of the juveniles reflect age-related differences in true survival rather than higher emigration of young birds, although we cannot completely rule out the latter possibility.

There are few other estimates of Twite survival against which to compare our results. Although measured over only three years, the survival of adults in this study (0.29 - 0.56) was slightly higher than that reported for populations in the South Pennines and N.W. Scotland (Table 3). By contrast, the survival of juveniles (0.14-0.34) was low relative to South Pennine birds. When compared to other seed-eating Cardueline finches, survival of adult Twite was similar to that for Linnet *L. cannabina* and Goldfinch *Carduelis carduelis* in the UK whereas survival of juveniles was notably poorer than for these species (Table 3). This suggests that low juvenile survival may be an important demographic driver of Twite population decline. One possible cause is the food availability over winter, since Atkinson (1998) found that the distribution of Twite wintering on salt marshes in late winter/early spring was strongly correlated with the density of Glasswort *Salicornia* seeds, and it is possible that through inter- and intra-specific competition the juveniles are more affected by food scarcity, especially as these habitats are eroding rapidly (Hughes & Paramor, 2004). However, it is also possible, although perhaps less likely, that mortality of first-year birds was higher during the breeding season. Additionally, estimates of survival from ring recoveries used by Siriwardena *et al.* (1998) are not as strongly confounded by emigration, which could explain the lower juvenile survival found in our study.

We did not monitor changes in breeding productivity during this study, and thus cannot rule out the possibility that poor productivity has also contributed to the decline of the Derbyshire Twite population. Potential causes of low productivity include predation, poor weather, and a shortage of suitable seed during the breeding period (Raine 2006, Wilkinson & Wilson 2010), although the latter is less likely given the season-long provision of supplementary food. An additional potential threat, unique to this population, is disturbance at the nest sites, which are primarily sited inside a working quarry. Knowledge of Twite nest use of the quarry is poor and warrants further work to identify nest sites and quantify causes of nest failure.

The decline of the English breeding population of Twite has been attributed to habitat deterioration on both the breeding and wintering grounds (Atkinson 1998, Raine 2006, Brown & Grice 2005), but the relative importance of these drivers is unknown. Conservation action to date has focused on improving breeding habitat conditions in the core South Pennines range, with conservation measures to increase summer seed food sources and protect nesting habitat. However, given the low apparent survival of juveniles recorded by both this study and Wilkinson *et al.* (in prep) for birds from different parts of the wider South Pennines breeding distribution but which share the same east coast wintering areas (Dunning *et al.* 2016), we suggest that conservation efforts are directed at improving the survival of Twite during their first year. Furthermore, given the scale of the decline in the South Pennines population (Wilkinson *et al.* 2018), this work should be coupled with monitored trial interventions (Christie *et al.* 2019) to enhance food availability during the non-breeding period, perhaps using supplementary seed in the short-term, at sites holding Twite in late winter.

## Acknowledgments

We would like to thank Steve Christmas (of Christmas & Christmas ringing group) for access to data and useful comments on early drafts of this manuscript. We would also like to thank Katrina Aspin, the Derbyshire Ornithological Society (DOS), RSPB, British Birds Charitable Trust, and CEMEX who have provided additional support for the English Twite monitoring project.

